# Combining phenotypic and genomic data to improve prediction of binary traits

**DOI:** 10.1101/2022.08.30.505948

**Authors:** Diego Jarquin, Arkaprava Roy, Bertrand Clarke, Subhashis Ghosal

## Abstract

Plant breeders want to develop cultivars that outperform existing genotypes. Some characteristics (here ‘main traits’) of these cultivars are categorical and difficult to measure directly. It is important to predict the main trait of newly developed genotypes accurately. In addition to marker data, breeding programs often have information on secondary traits (or ‘phenotypes’) that are easy to measure. Our goal is to improve prediction of main traits with interpretable relations by combining the two data types using variable selection techniques. However, the genomic characteristics can overwhelm the set of secondary traits, so a standard technique may fail to select any phenotypic variables. We develop a new statistical technique that ensures appropriate representation from both the secondary traits and the phenotypic variables for optimal prediction. When two data types (markers and secondary traits) are available, we achieve improved prediction of a binary trait by two steps that are designed to ensure that a significant intrinsic effect of a phenotype is incorporated in the relation before accounting for extra effects of genotypes. First, we sparsely regress the secondary traits on the markers and replace the secondary traits by their residuals to obtain the effects of phenotypic variables as adjusted by the genotypic variables. Then, we develop a sparse logistic classifier using the markers and residuals so that the adjusted phenotypes may be selected first to avoid being overwhelmed by the genotypes due to their numerical advantage. This classifier uses forward selection aided by a penalty term and can be computed effectively by a technique called the one-pass method. It compares favorably with other classifiers on simulated and real data.

## 1. INTRODUCTION

Plant breeders typically select a desirable trait such as yield and then cross genotypes that have the highest yield in the hope of increasing genetic gains. After crosses are done, the number of genotypes can be infeasibly large to screen in fields. There are many properties that ideal genotypes should have: resistance to drought, resistance to salinity, minimum days to 50% maturity, height, etc. Some of these properties are categorical; continuous properties can be discretized and then regarded as categorical as well. The problem then becomes how to identify, effectively, the genotypes that are most worth testing in fields. For this identification, we can use predictive models to select untested genotypes for the main trait of interest based on their marker profiles and their associated traits. In many cases, the associated traits are easier to measure than the main trait because they do not require extensive field testing. More formally, the problem is, given the availability of marker data, how to leverage the information from the associated traits to improve predictions. This problem is even more difficult when the data types for the main trait have very different dimensions.

Suppose we have *p* associated traits that we call ‘macroscopic’ or phenotypic explanatory variables *U* = (*U*_1_, …, *U*_*p*_)^*T*^ and we have *q* ‘microscopic’ or genomic variables, *W* = (*W*_1_, …, *W*_*q*_)^*T*^. Single Nucleotide Polymorphisms (SNPs), gene expression data, and sequencing data are amongst the most common. We focus here on agronomic data but this situation often arises with other biomedical data where the macroscopic variables may correspond to demographic status, health history, or test results. In such a ‘multi-type data’ scenario, commonly the microscopic variables massively outnumber the macroscopic variables. The task is to form a classifier that makes use of both the phenotypic and genomic data.

Suppose a plant is in class *Y* = *c*, for some *c* ∈ {1, …, *C*}. One simple technique for developing a classifier *Ŷ* that predicts *Y* is to concatenate the explanatory variables into a single *p* + *q* vector (*U*_1_, …, *U*_*p*_, *W*_1_, …, *W*_*q*_)^*T*^ and then apply any decent classification technique to form *Ŷ*. However, this may give poor results because, for *q* ≫ *p*, the genomic variables will outcompete the phenotypic variables to explain *Y*. Few, perhaps none, of the phenotypic variables will be retained even when it is implausible for all of them to be unimportant. In effect, the genomic variables are so numerous that spurious relationships between *Y* and *W* ‘crowd out’ the explanatory variables in *U*.

This is a common phenomenon, most easily seen in linear regression where *Y* is continuous. Suppose there are *n* observations *y* = (*y*_1_, …, *y*_*n*_) and the phenotypic and genomic data are combined into one design matrix *D*. Then the fitted values are *Ŷ* = *D*(*D*^*T*^ *D*)^−1^*D*^*T*^ *y*, provided the inverse exists or a generalized inverse such as Moore-Penrose is used. The matrix *D* is *n* × (*p* + *q*) so, when *n ≤ p* + *q, y* can be exactly represented by many possible linear combinations of the explanatory variables. Moreover, when *q ≫ n* and *n > p*, most of the linear combinations that express *y* perfectly will involve the genomic variables only thereby leaving out the phenotypic variables for mathematical reasons and giving poor predictions. The same principle carries over to classification.

A multiclass setting – *C* ≥ 3 – suggests that we should use a multiclass classifier. However, the consensus view seems to be that the performance of multiple binary classifiers is no worse and often better than using a single multiclass classifier; see Rifkin and Klautau (2004), Sánchez-Maroño et al. (2010), and Hosenie et al. (2019). The situation for plant breeding is potentially more complicated than just having many classes because traits may work against each other. For instance, yield and protein content are complex traits affected by a large number of gene effects, and may be inversely related. For this reason, there may be numerous categories and it will be important to have a single good classifier that can accommodate a large number of classes to help determine which of many crosses are most likely to result in cultivars with the optimal traits.

The method we propose here can be regarded as an elaboration of the approach taken in other settings in what is often called genomic selection. For instance, the review by Desta and Ortiz (2020) uses the same sort of gene × environment interaction interpretation – essentially a components of variance model – but restricted to single genomic data types. This class of models has been used in plant and animal breeding typically using genetic markers. The key examples in their review differ from our methodology mainly because we use two data types and must account for the possibility they contain some information in common about the main trait. However, Desta and Ortiz (2020) includes considerations of trait complexity and this leads them to argue that nonlinear models will, as a rule, give better predictions than penalized linear models. Our results are consistent with this intuition.

Earlier efforts to predict genomic breeding values in animal science, specifically dairy bulls, have long been modeled as traditional regression problems using linear and nonlinear techniques. One example is Moser et al. (2009) who regressed breeding values on SNP marker and pedigree data. In this work, fixed effects linear regressions were found to underperform the other four methods tested, but these other four techniques performed similarly well predictively. These four techniques represented conceptually disjoint methodologies – none of which were linear in the conventional sense. The authors noted that combining two data types gave better results than either single data type although the degree of improvement depended on the trait studied. They avoided the problem of different dimensionality of the data types by dimension reduction that depended on all the variables as in partial least squares (that can mix explanatory and response variables), kernel methods, a form of Bayes regression on all variables, and ridge regression. These techniques often give little sparsity.

More recently, Ma et al. (2018) has used genomic data to predict phenotypes in a species of wheat, another example of the class of problem we address here. Like us, they argue that ‘genomic selection has the inherent advantages of predicting phenotypic trait values of individuals before planting’. Rather than seeking sparsity, they used a model average of a deep convolutional neural network and essentially a random effects model showing that the model average outperformed its two components as well as several other related techniques. One possible drawback of this method is its complexity. However, there is evidence of a principle of matching between the complexity of data and the complexity of techniques used to model it. So, such complexity may often be unavoidable. One of the original motivations for neural networks was the hope that they could accommodate multitype data and while Ma et al. (2018) did not investigate this, neural networks remain a promising approach for genomic selection of complex traits with multitype data.

Another methodology for genomic selection using genomic data is given in Jeong et al. (2020). Recognizing that genomic data is usually high dimensional they developed a technique for doing variable selection first – hoping to find the optimal genomic variables, often taken to be SNPs – before making predictions. They use a mixture of *p*-values and correlations to select which markers to include in their predictors which are from machine learning models. Thus they seek a double sparsity from variable selection combined with the natural sparsity of methods such as random forests or penalized neural networks. While these authors only use genomic data from soybeans and rice, their techniques generalize to other settings and extend to allow interactions between genomic variables, a more difficult problem. Much of the success of their technique probably rests on good variable selection in very high dimensions for variables that are broadly comparable.

From this it can be seen that there are numerous approaches to the genomic selection problem or more precisely the problem of predicting phenotypes from other data types. Our contribution is an explicit treatment of the use of two data types (one high dimensional and one low dimensional) to construct sparse binary classifiers that retain variables from each data type and, sometimes only incidentally, provide sparsity that can be useful for understanding underlying mechanisms. Our classifier avoids the ‘crowding out’ problem and is relatively simple in structure and implementation even as it allows for complex traits. It also outperforms many other classification techniques in terms of misclassification rate. We have chosen logistic classifiers because they are more interpretable than most other methods, often more stable, and relatively well understood.

In Sec. 2 we give an informal presentation of our methodology to highlight the intuition guiding it. In Sec. 3 we provide the mathematical details of our methodology, including our computational procedure. In Sec. 4, we compare our method to a wide variety of other techniques on simulated data intended to be similar to the real data from maize breeding that we analyze in Sec. 5. In Sec. 6 we look at another real data set using soybeans where our general method is equivalent to a simplified version of it in the special case that the secondary traits are highly correlated. Finally, we discuss the implications of our findings in Sec. 7 and briefly summarize our results in Sec. 8.

## 2. OVERVIEW OF THE METHODOLOGY

The shortcoming of standard penalized logistic regression high-dimensional classifiers in our context is that they do not have a way to include phenotypes in the classifier first to avoid them being overwhelmed by the genomic variables. This leads us to a forward selection approach to include variables one by one, where it is easy to accommodate the former type first. However, forward selection relies heavily on a stopping rule. The reliance can be reduced by using a penalty term at each iteration of forward selection as in Hwang et al. (2009). This can filters out insignificant predictors and control model size. Another issue is that the phenotypes may already contain some effects of the genomic variables, so in the first stage of modeling, only intrinsic effects of phenotypes should be counted. This necessitates a step to separate the effects of genomic variables from the phenotypic ones before the latter are selected. This ensures that a potential effect of a genomic variable is not wrongly counted as the effect of a phenotypic variable in the classification rule.

We construct our binary logistic classifier in two steps. First, to extract the intrinsic effects of *p* phenotypic variables (*U*_1_, …, *U*_*p*_), we regress each of them on the set of genomic variables (*W*_1_, …, *W*_*q*_) and collect the regression residuals. However, as each regression is very high dimensional, an ordinary least square regression is unstable. A penalized regression, such as the least absolute shrinkage and selection operator (LASSO) (cf. Tibshirani (1996)) allows a more stable estimation with interpretation. To handle high correlations among the explanatory variables, we actually consider some variations of the LASSO such as the adaptive LASSO (Zou (2006)), the adaptive elastic net (AEN) (Zou and Zhang (2005)), and the smoothly clipped absolute deviation (SCAD) (Fan and Li (2001)). We then collect the regression residuals, which we denote by (*V*_1_, …, *V*_*p*_), and let them replace (*U*_1_, …, *U*_*p*_). We use sparse regressions of the macroscopic variables on the microscopic variables in this step for the same reason as noted in Sec. 1: If we don’t then we risk simply representing them perfectly by non-unique linear combinations of the microscopic variables. We comment that our motivation is is to adjust the phenotypic variables by sparse regression on genotypes so as to extract their intrinsic information. However, the more accurate way to regard this step as an adjustment to them. This will be seen more clearly in Sec. 6.

Second, we use a sparse logistic regression of *Y* on (*V*_1_, …, *V*_*p*_, *W*_1_, …, *W*_*q*_) by a penalized forward selection method including potential *V* s first before allowing any *W*s. We call this the one-pass method (OPM) because we allow only the most significant predictor to enter at each stage in forming the logistic regression rule and obtain the coefficients by independent Newton-Raphson updates. This approach worked well in other contexts (see Hwang et al. (2009), Ghosal et al. (2016), Turnbull et al. (2013), Luo and Ghosal (2015)) and a convergence theorem is provided by Luo and Ghoshal (2016). The ‘trick’ that makes our method successful in the multitype data setup with a severe unbalance in the values of *p* and *q* is that we order the *p* + *q* variables for inclusion (macro first, micro second) and then consider the use of the adjusted macro/phenotypic variables in the model first. This approach allows us to build a sparse classification rule that includes the relevant phenotypic variables. Phenotypic variables can be readily interpreted and their interpretation may be augmented by a relatively small number of micro/genomic variables. These latter variables may be few enough that they, too, become interpretable.

The second step has a key feature of our approach: We have not done an optimization over the entire parameter vector as in a GLMNET; see Friedman, Hastie, and Tibshirani (Friedman et al.).

We use forward selection, one variable at a time, with an objective function that includes a penalty term to eliminate irrelevant variables more effectively. Since the forward selection optimization is performed one at a time, high dimensional data sets can be handled by parallelizing the procedure. Penalized forward selection can find sparse models effectively without losing predictive power.

## 3. FORMAL PRESENTATION OF THE METHODOLOGY

Suppose we have data points denoted *y*_*i*_, *u*_*i*_ = (*u*_*i*1_, …, *u*_*ip*_), and *w*_*i*_ = (*w*_*i*1_, …, *w*_*iq*_) for *i* = 1, …, *n*, and that the *u*_*ij*_s for *j* = 1, …, *p* and the *w*_*ik*_s for *k* = 1, …, *q* are entries in vectors of *n* outcomes of *U*_*j*_ and *W*_*j*_ respectively. Without loss of generality, we assume E(*U*) = 0, E(*W*) = 0, Var(*U*) = 1_*p*_, and Var(*W*) = 1_*q*_, respectively *p* and *q*-dimensional identity matrices. In practice, this is enforced by studentization, i.e., replacing *u*_*ij*_ by 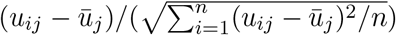, where 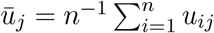 and treating the *w*_*ij*_s similarly.

Beginning with Step 1, for each fixed *j*, the *u*_*ij*_ values can be regressed on the *w*_*i*_ = (*w*_*i*1_, …, *w*_*iq*_) in a linear regression model

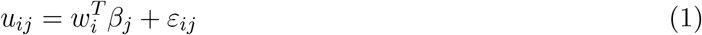

where *β*_*j*_ = (*β*_*j*1_, …, *β*_*jq*_)^*T*^ and *ε*_*ij*_ is a vector of *n* mean-zero finite variance errors. Then each *u*_*ij*_ is replaced by its regression-residual 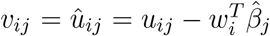, where the regression coefficients 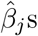 s are found by minimizing the sum of squares of residuals with a penalty term

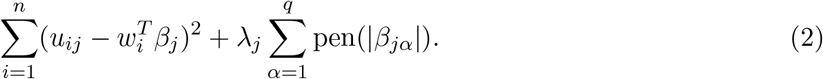

Here, the function pen : [0, ∞) → [0, ∞) can be taken to be the LASSO penalty (absolute value) or one of its variants like the ALASSO, AEN, or SCAD penalty. In computational work to be presented shortly, we found that AEN gave the best results. We suggest that this is the case because the AEN penalty is a combination of the ridge regression penalty that is not sparse and the LASSO penalty that does shrinkage and selection at the same time and hence does give sparsity. Since our methodology is proposed for data that has some sparsity but is not truly ‘sparse’ the form of the AEN penalty may capture something intrinsic about the data.

Now, for Step 2, suppose that the explanatory variables are written as *z*_*i*_ = (*z*_*i*1_, …, *z*_*i,r*_) = (*v*_*i*_, *w*_*i*_) = (*v*_*i*1_, …, *v*_*ip*_, *w*_*i*1_, …, *w*_*iq*_) for *i* = 1, …, *n*, and *r* = *p* + *q*. The task is to use the *y*_*i*_s and *z*_*i*_s to develop a classifier for future outcomes of *Y*. We control the order of entry of the *z*_*i*_s into the classifier, starting with the residuals from the macrovariables and applying the OPM to each of them in order. This ensures they will not be overwhelmed by the genetic variables. Let Γ = ((*γ*_*cj*_)) be a *C* × *r*-matrix of coefficients defining the multinomial logistic model having *C* cells. We allow an arbitrary positive integer *C* ≥ 2 although we illustrate the binary (*C* = 2) case only. The probability mass function is

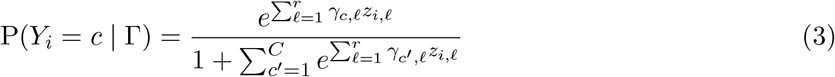

and our classifier for the *n* + 1 data point is

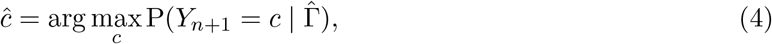

where 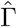 is an estimate of Γ using *D* = {(*y*_1_, *z*_1_), …, (*y*_*n*_, *z*_*n*_)}.

Our basic approach to estimating Γ will be a Newton-Raphson method applied the likelihood penalized by a LASSO term; other penalties such as SCAD and AEN did not give as good results.

The problem is that *r* = *p* + *q ≫ n* and *q ≫ p*; this will necessitate a sparsity method on Γ (in addition to that on (2)), proper selection of the *γ*_*c,ℓ*_s to update, as well as the more familiar updating expressions that will require ‘learning rates’ *λ*_*c,ℓ*_ and a stopping rule appropriate to the selection of the *γ*_*c,ℓ*_s and *λ*_*c,ℓ*_s. Note that for the two adaptive penalized methods, AEN and ALASSO (but not SCAD), we must specify a *c* × *r* matrix of *λ*_*c,ℓ*_s, analogous to the *λ*_*j*_s in (2).

### 3.1 Setting up a Newton-Raphson (NR) method

To find values for the *γ*_*c,ℓ*_s, we require the first two derivatives of the likelihood. So, consider a parameter *γ*_*c,ℓ*_ for some fixed values *c* and *ℓ*. As a function of *γ*_*c,ℓ*_, the log-likelihood (3) is

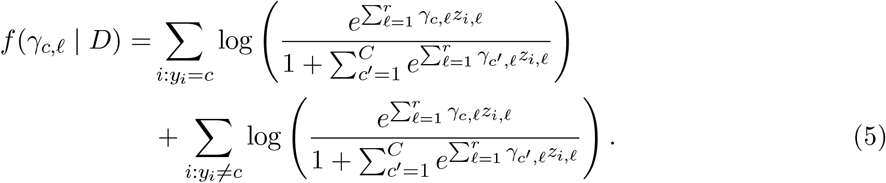

The maximizer of (5) is found by a simple modification of the NR method, which requires the first two derivatives of (5), but in the presence of the penalty term (2), we need to maximize

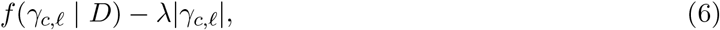

with respect to *γ*_*c,ℓ*_. This optimization problem has been solved in Wu et al. (2009) and Chap. 2.3 of Burden and Faires (2011). Let

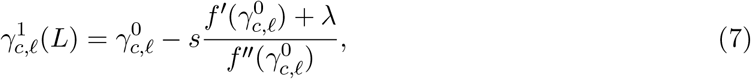

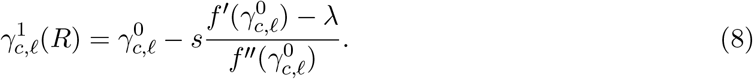

respectively corresponding to the left- and right-derivatives of (6) with respect to *γ*_*c,ℓ*_. If 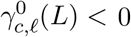, set 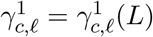 and if 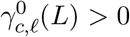, set 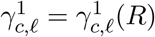. If either *γ*_*c,ℓ*_(*L*) = 0 or *γ*_*c,ℓ*_(*R*) = 0, then set 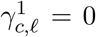. The update of 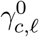to 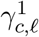will always be unique (because *f*^*″*^ (*γ*_*c,ℓ*_) *<* 0 for all *γ*_*c,ℓ*_, unless all *z*_*i,ℓ*_ = 0) and *λ* must be chosen uniformly over all the *c*’s and *ℓ*’s. The same updating procedure generates 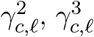 etc. Iterations of this procedure are used in the algorithm of Subsec. 3.2 until convergence is achieved.

The extra factor *s >* 0 in (7) and (8) is a step length, usually *s ≤* 1. Dennis and Schnabel (1996) showed several examples where the optimizations failed when the step length parameter was fixed at 1. Thus, line search methods such as step halving Schoenberg (2001), golden section Gill et al. (2019) are usually preferred to select the step length parameter optimally at each iteration. Here we have used the step halving strategy.

Procedures analogous to the above for LASSO were used for AEN, SCAD, and the raw data. The results are not included here because these shrinkage methods clearly underperformed LASSO. We surmise from this that the use of a sparse model from doing selection and shrinkage simultaneously as is done by LASSO helps give a single sparse model and the sparsity is more important in the OPM than preserving all variables that might be relevant.

### 3.2 Algorithm

Since the explanatory variables are studentized, we initialize by setting *γ*_*c,ℓ*_ = 0 for all *c* and *ℓ*. Let PLL(*c, ℓ*) stand for the penalized (*c, ℓ*)-th (univariate) log-likelihood. To minimize the PLLs, write the *m*-th update of Γ as 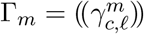 so Γ_0_ = Γ. Then

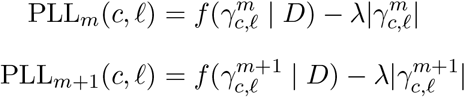

are the ‘old’ and ‘new’ values of the PLL occurring in the NR procedure.

Steps in the OPM algorithm The basic idea is to update all the *γ*_*c,ℓ*_s that are zero up to a certain tolerance and then choose the (*c, ℓ*) that gives the largest value of PLL_new_(*c, ℓ*). Then, if *γ*_*c,ℓ*_ ≠ 0, update it and cycle through all the other *γ*_*c,ℓ*_s, stopping when the largest value of PLL_new_(*c, ℓ*) occurs for some *γ*_*c,ℓ*_ = 0. Formally, let *ϵ >* 0 define the tolerance on the NR convergence and assume that Γ_0_ is the result for *m* = 0. Now, the OPM, i.e., the univariate NR updating starting from stage *m*, is as follows.

1. Update each 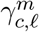 that is zero by using the (univariate) NR method, i.e., (7) and (8). The iterations continue until two criteria are met. First, is the tolerance criterion

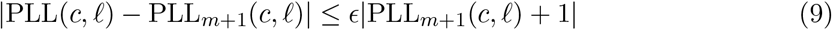

depending on *ϵ*; second is the minimality criterion

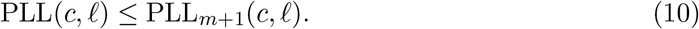

Both should generally be strict inequalities, and neither should be violated in a valid update. The NR updates for (*c, ℓ*) begin with *s* = 1. If the resulting 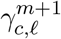 do not satisfy (9) and (10), the procedure is repeated using *s* = 1*/*2, 1*/*2^2^, 1*/*2^4^, …, 1*/*2^10^. If strict inequality in one of (9) and (10) is achieved with only in the other, the update is also accepted. If the search over step sizes does not yield an improvement, set 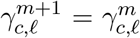. Write PLL_*m*+1_(*c, ℓ*) for the final value of PLL from the univariate NR updates of PLL_*m*_(*c, ℓ*).
2. Given all PLL_*m*+1_(*c, ℓ*)s, let

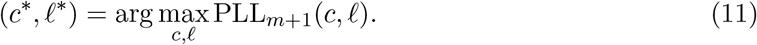

If 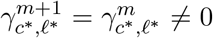 then the updating is zero and the parameter value is not zero. So, set Γ^*m*+1^ = Γ^*m*^; the final estimate of Γ is then Γ^*m*^. If 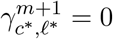, then no variable is selected and again the final estimate of Γ is Γ^*m*^. If 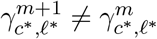, then repeat Step 1.
3. Stop when no variable is selected, i.e., the largest values of the univariate PLLs only occur for (*c, ℓ*) that give *γ*_*c,ℓ*_ = 0.

### 3.3 Tuning parameters

The tuning parameters *s, ϵ* and *λ* must be chosen appropriately. Good values for *s* are already built into the algorithm. The value of *ϵ* must be chosen small enough that (9) can be achieved with reasonable run-time of the OPM. For the simulations reported in Sec. 4, we used *ϵ* = 10^−4^. They are based on approximately 5000 explanatory variables somewhat mimicking our real data. Larger values like 10^−3^ seem to blur the comparison between the methods while smaller values like 10^−5^ lead to excessive computational cost without noticeable benefits.

Since *λ >* 0 is unidimensional, cross-validation (CV) is a good default as will be seen in the examples. We then used a simple grid search on a range of *λ*s. In each case, the class predictor in is used and compared to the actual class via misclassification error.

### 3.4 Convergence properties of the algorithm

Observe that each univariate log-likelihood is strictly concave and the penalty is concave, although not strictly. Nevertheless, the univariate PLLs are sums of a strictly concave and a concave function. Thus, each univariate PLL is strictly concave. Moreover, an important aspect of this method is that for a given *ϵ >* 0, the convergence is guaranteed within a fixed number of steps by Kantorovich’s theorem for the NR algorithm; see Gálantai (2000). This ensures that the objective function increases over iterations.

The algorithm is initialized at the matrix Γ_0_ = 0 and the value of any 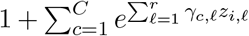 is (*C* + 1) at initialization. Therefore the minimum of the log-likelihood starts at −*n*(*C* + 1) ln(*C* + 1). The log-likelihood will always be negative because the likelihood is obtained from the joint probabilities of a multinomial distribution. In addition, the likelihood will increase by at least *ϵ*, after each iteration. Hence, the maximum number of steps to converge will be [*n*(*C* +1) ln(*C* +1)]*/ϵ*.

Methodologically, we find that sparse regression using LASSO, followed by the logistic classifier version of OPM penalized with AEN performs better than, or at least only slightly worse, than any competing method, for combining low and high dimensional explanatory variables for the purpose of binary classifiaction. In cases where our method is slightly numerically worse, around 1%, the interpretablity our method in terms of sparsity and variable classes more than compensates when all sources of error are considered.

## 4. SIMULATIONS

We use the R package `synbreed` to generate artificial data of the sort our method is designed to analyze. Given this, we compare the classification results from four versions of our method (based on different penalties) to the same four versions coded in the R package `BGLR`, the most common alternative to our procedure. We also compare our method to four standard machine learning techniques, Fisher’s linear discriminant analysis (FDA), support vector machines (SVM), random forests (RF), and boosting (for binary classification)

The first step is to generate data from `synbreed` and then transform it to provide a concatenated vector of relatively few macroscopic variables and relatively many microscopic variables. The next step is to generate our classifiers: i) four based on different shrinkage methods, logistic regression, and OPM; ii) four based on different shrinkage methods and `BGLR`, and iii) four standard machine learning methods. For the machine learning methods we use the data ‘as is’. That is, we do no pre-processing of the data e.g., we do not adjust the macro variables by a sprase regression. This means we obtain a fair comparison of the performances of the procedures. We next review how the data was generated; although some of the methodology is standard, we include these explanations for the sake of completeness.

To generate data using `synbreed`, we choose a number of chromosomes and their lengths. We choose ten because maize has ten chromosomes. To be specific we choose lengths (2.86, 1.83, 2.09, 1.89, 1.73, 1.38, 1.58, Gb, where Gb indicates gigabases i.e., 10^9^ nucleotides. The individual lengths are set according to the Genetic 2008 composite map found on the Maize Genetics and Genomics Database; see www.maizegdb.org. The total length is 17.97 Mb.

Next, `synbreed` internally generates two model genotypes that we choose to be 5000 markers each. These are the ‘parents’ assumed to be homozygous in all 5000 markers, i.e., we have two vectors of length 5000 with all entries 0 or 2. `synbreed` then creates 3500 progeny from the two founding parents by choosing one of the two values in each of the 5000 entries. This generates a 3500 × 5000 matrix of 0s and 2s, say *X*_*m*_ (*m* for ‘micro’) in which *n* = 3500 is the sample size and each entry in *X*_*m*_ is the number of copies of a nucleotide at a given location for a given subject, i.e., is 0 or 2. That is, given the number of chromosomes and their lengths, `synbreed` generates *X*_*m*_ and this process is valid for all double haploids such as maize. For more details on how `synbreed` generates synthetic data via a random-effects model, see Wimmer et al. (2012) and Wimmer et al. (2012). To finish the generation of the full simulated data set we randomly pick 20 columns of *X*_*m*_ to represent quantitative trait loci (QTLs). This means that the other 4980 columns of *X*_*m*_ are markers that, *in principle*, should have no effect on our classification apart from the linkage disequilibrium (LD) considerations discussed below.

We generated 3500 samples so that later we can subsample from them. As will be seen, this lets us ensure the correct proportions of 0s and 1s in each of the 20 repetitions of the subsampling. For each subsample of size 277 (the sample size in our real data set) we sample genotypes to ensure a bimodal distribution with the desired proportions for each group. This is necessary in our framework because we simulate the main trait first as a continuous variable then we use the mean as threshold to form groups 0 and 1 thereby giving a bimodal distribution based on the proportions of 0s and 1s. The details are given in Subsec. 4.3. An alternative would be to generate repeated samples of size, say, 277. In this case, however, it is not clear how the desired proportions of 0s and 1s could be ensured.

Next, we describe how the output of `synbreed` can be manipulated to get data in the form we want for outcomes of a ‘main’ trait, say *Y*. In the subsection after that, we repeat this procedure to generate macroscopic variables to be selected from to form classifiers.

### 4.1 Generating the main trait

Consider the expression

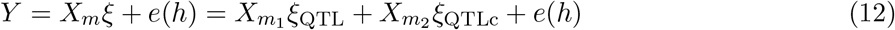

in which *Y* is the response variable that interests us, e.g., yield, *ξ* is a real 5000-dimensional random vector representing the effect of the markers in *X*_*m*_ and *e*(*h*) is an independent error term that affects the heritability *h ∈* (0, 1) of the QTLs. On the right side of (12), the regression function is split into two terms, also assumed to be independent. We write 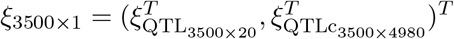 and 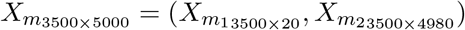, where QTLc means the complement of the QTLs and the dimensions of the vectors are indicated by sthe numbers. The 3500 × 20 matrix 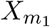 represents the columns of *X*_*m*_ corresponding to the 20 QTLs and the 3500 × 4980 matrix 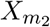 represents the portion of *X*_*m*_ corresponding to the null effect markers. Their coefficients are *ξ*_QTL_ and *ξ*_QTLc_, respectively. These expressions for *ξ* and *X*_*m*_ are ‘conceptual’ because in reality which marker effects should be part of the main trait 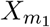 *ξ*_QTL_ are unknown.

The heritability *h*^2^ is the proportion of the variability of *Y* explained by the genetic components:

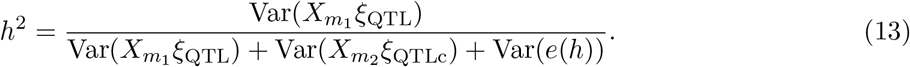

To simulate from (12), start by generating the 20 entries in *ξ*_QTL_ independently from a N(0, 5^2^). Then, *X*_*m*1_*ξ*_QTL_ is the ‘signal’ portion of *Y* also called the genetic estimated breeding values, GEBV. If 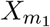 is taken as fixed it is not hard to find Var(GEBV). If 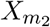 is fixed as well, the ‘non-signal’ portion of *Y* is NS 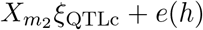. In reality, typically 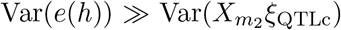 whereas in (12) or (13) we usually have 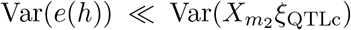 because the number of non-QTLs is very large even though each effect may be very small.

We can now imagine one of two scenarios. The first is that 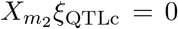 exactly and we construct the error term. The other is that we set *e*(*h*) = 0 and construct a viable 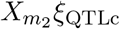 for (13). The difference is whether the variability in the main trait *Y* is assumed to arise from measuring *Y* or from the QTLcs that contribute variability to *Y* but do not affect its location. We choose the latter and hence set *e*(*h*) = 0 because generating *e*(*h*) via independent normals, would only make any patterns harder to identify.

Suppose *h*^2^ = 0.5, a typical value, and that *ξ*_QTLc_ is generated from 4980 independent N(0, *ν*^2^*/*4980) random variables, i.e., all the complementary QTLs are the same in terms of their distribution, where 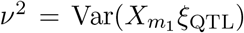. Then, since a half of the phenotypic variability is due to genetic factors, 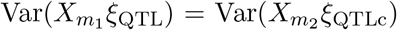, by taking the term-wise expectation of (12). Set 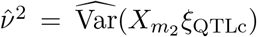 and define 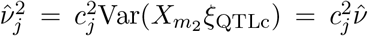, where *c*_*j*_ ranges over {0.01, 0.02, …, 0.99} for 1 *≤ j ≤* 99. Set 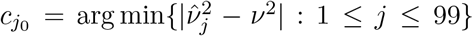 and replace 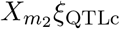 by 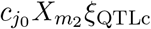, i.e., rescale (down or up) 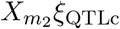 so that we get heritability 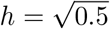 of the signal 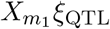 with respect to the total variability. Then (13) becomes

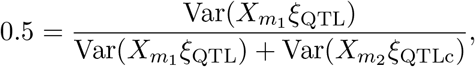

giving 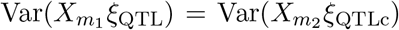 i.e., the non-signal portion of the main effect is half of the signal and 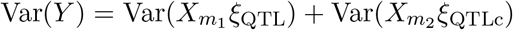 so that 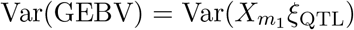. To enforce the heritability constraint *h*^2^ = 0.5, we generate *Y* by multiplying 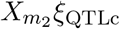 by 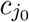 (a scaling constant) and write the 3500 × 1-vector of responses as

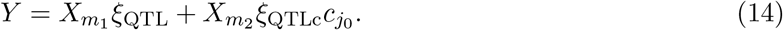

In the alternative version where we set 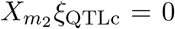 we are using the values in 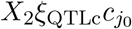 as approximate outcomes of the random variable *E*(.5) (or more generally *NS*). In a strictly heuristic sense, 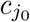 is summarizing the effect of the heritability *h* in (12).

### 4.2 Generating macroscopic variables

Next, we modify the procedure in Subsec. 4.1 to generate outcomes of the macroscopic traits. Suppose we want to generate *p* = 20 macroscopic traits for the same 3500 ‘subjects’ as before. These can be constructed from the matrix *X*_*m*_ within `synbreed` by inputting distances and directions. For simplicity, we choose *±*1, …, *±*10, that is, the first macroscopic variable will be constructed by modifying the location of the first of the 20 QTLs in Subsec. 4.1 by moving it +1 from its actual location. The effect is to make the output for the first QTL slightly different from the original QTL. The reason the new QTL ‘+1’ is different from the old QTL is linkage disequilibrium (LD): The effect of the marker slightly to the right or left of the original marker is slightly different from the original marker. The linkage between the QTL effect and its location implies that no linear function of the new QTL’s will replicate the main trait. (The term disequilibrium means that moving the location of a marker by a small amount does not make the marker effect go to zero.) Within `synbreed`, the first macroscopic trait is selected randomly from the original 20 QTLs used to find the main trait *Y* (modified by LD using the vector of distances and directions). For example, if 15 of the 20 are chosen, the actual form of the first macroscopic variable is

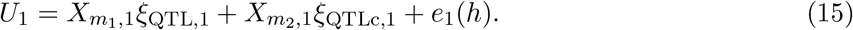

The variables in (15) are the analogs of the variables in (12). In particular, *U*_1_ is a 3500 × 1 vector that will be modified in the same way that *Y* in (12) was modified to give the *Y* in (14), 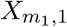 is the 3500 × 15 submatrix of *X*_*m*_ corresponding to the 15 QTLs (as modified by LD), *ξ*_QTL,1_ is a 15 × 1 vector of marker effects for them, 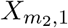 is a 3500 × 4985 submatrix of *X*_*m*_ corresponding to the other 4985 markers (modified by LD), *ξ*_QTLc,1_ is the 4985 × 1 vector of the markers not included amongst the QTLs, and *e*(*h*) is the error term depending on the heritability *h*. Following the procedure in Subsec. 4.1, we modify *U*_1_ by imposing *h*^2^ = .5. After subtracting residuals from the sparse regression in (2), we have *V*_1_. This procedure can be repeated to generate *V*_2_, … *V*_20_. Within `synbreed`, the random selection of QTLs is done so that the number of QTLs in each iteration is nonincreasing and the locations of the traits are shifted according to the next distance/direction to enforce LD. This gives *U*_2_ as an analog to *U*_1_ from (15). So, the procedure in Subsec. 4.1 can then be used to generate *V*_2_, and analogously *V*_3_, …, *V*_20_. Without loss of generality, we assume the data points for each *V*_*j*_ are standardized.

The result is that we now have the main trait *Y*, the vector *V* of macroscopic variables, and our original 3500 × 5000 matrix *X*_*m*_. Clearly, *X*_*m*_ corresponds to the matrix *W* of standardized *W*_*i*_s (see (1) with *q* = 5000). Since the *V*_*i*_’s are standardized as well, the *Z*_*i*_s are standardized and we assume they can be treated as continuous.

### 4.3 Comparisons on simulated data

Recall that the procedure described here involves two basic steps. In the first, two types of data vectors *U* of length *p* and *W* of length *q* where *p ≫ q* are used. The point is to regress each of the *p* variables on the *q* variables sparsely and then replace *U* by its residuals, say *V*. The second stage is to form the variable *Z* = (*V, W*) of length *p* + *q* and sparsely regress the class indicator *Y* on *Z* for the logistic classifier. The resulting classifier is thus formed from *p* corrected ‘macroscopic’ or phenotypic variables and *q* ‘microscopic’ or genomic variables. Loosely, the information in *V* is the information in *U* that is not in *W*. Now, updating the *γ*_*c,ℓ*_s by using *V* before *W* should prevent the microscopic variables from spuriously overwhelming the macroscopic variables.

Given data simulated as described in Subsecs. 4.1 and 4.2 with *n* = 3500, *q* = 5000 and *p* = 20, we randomly select subsamples of size 277, the same size as the real data set we analyze in Sec. 5, following the scheme below that ensures fixed proportions of 0s and 1s. Then on each subsample, we compare our sparse regression-ordered-OPM under three different penalties (SCAD, ALASSO, AEN) with the analogous results under the standard R-package `BGLR` and with the results of four other standard classifiers, FDA, SVM, RF, and boosting (in the binary case). Implementation of these classifiers was done using the R-packages `PenalizedLDA` for FDA, `CMA` for SVM and RFs, and `maboosting` for boosting. In all cases, we use the training data to find empirical values of tuning parameters in the OPM and for the prior *π* in `BGLR`. When we used (2), the decay parameter was found internally to the packages. For comparison purposes, we also included what we call the ‘RAW’ method corresponding to feeding the macro and micro variables into the OPM. This uses no penalizations. It simply does independent Newton-Raphson approximations to find coefficients for the input variables, starting with the macro variables.

For each of the 20 replications, the construction of each sample of size 277 is done using proportions 0.5, 0.65, 0.75, 0.85, and 0.95 to define the classes i.e., the 0s and 1s. To do this, we randomly choose 20 subsets of size 277 from the 3500 data points, ensure they have the appropriate proportions of 0s and 1s, and then partition them into subsets for training and testing for each method. To ensure we have the correct proportions of 0s and 1s, we assign 0 to those of the 3500 that have value smaller than zero and 1 otherwise. Then we used the 25th percentile on the left and the 75th percentile on the right, and the corresponding SDs, to impose a distribution on each side. This means that few of the original data points near zero are selected. The rest are assigned 0 or 1 according to the population we want.

For the OPM-methods, we partition each sample of size 277 into three parts: 120 for estimating the *β*s, 80 for estimating *λ* in the shrinkage method, and 77 for testing. For the machine learning methods, we use a training set of size 200 and a test set of size 77. In all cases we found the (empirical) misclassification rate by the formula 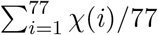 where *χ*(*i*) = 1 if the predicted and actual class of data point *i* are different i.e., we make an error, and *χ*(*i*) = 0 if they are the same, i.e., no error is made.

Different classifiers can perform differently for different proportions – one that performs relatively well at .5 may perform relatively poorly at .85. For instance at .99 a trivial classifier that chooses only class one will have an error rate of 1%. Accordingly, in our simulations, we searched over the range of proportions to understand how well the various classifiers performed. Since classification with a true proportion *α ∈* (0, 1) is equivalent to classification with 1 − *α ∈* (0, 1) it was enough to cover the range from .5 to 1. Moreover, there is no point using classification when *α* = 1 and for values of *α* close to one, all classifiers tend to ‘bail-out’: No matter what class a subject is in, the classifier simply returns the biggest class because, past a certain *α*_0_, the error of misclassification using the biggest class is smaller than the error from trying to identify a subject’s class nontrivially. We found that bailout generally occurred for *α ≈* .95, not an unusual finding. Hence, we chose .95 as the highest proportion for computation to verify that we saw bail out. Often values of *α* near .5 give fairly stable performance so we chose a slightly larger distance between .5 and the next higher proportion, .65, than between the other proportions which differed by .1 simply as way to cover the range.

A separate consideration is that, in the five real data cases that we used, (namely Pan161, Pan162, Pan163, Pan169, and Pan171 in the notation of Liang et al. (2020)) the true values of the proportions were approximately .59, .65, .74, .95, and .59, respectively. Thus, values around .65 were most representative for our work in Sec. 5.

The methods we use can be segregated into three different classes. First, there are the ‘doubly sparse’ methods such as we advocate here – few terms and few variables. Second, there are singly sparse methods such as SVMs that have few terms but many variables. Finally, there are non-sparse methods that permit many terms and many variables such as RF, boosting, and FDA. It has been suggested that RFs often have some sparsity properties in practice e.g.,there are relatively few important variables, but this is not built into the methodology and, indeed, we did not find that here. On the other hand, RFs often work inexplicably well. Intuitively, one expects that non-sparse methods should work best with non-sparse data, single sparse methods should work best when the data have some sparsity, and doubly sparse methods should work best when the data really is sparse.

Altogether, we used four versions of the OPM (ALASSO, SCAD, AEN, and RAW), four methods from `BGLR` (ALASSO, SCAD, AEN, and RAW), and four machine learning methods (RF, boosting, SVMs, and FDA) each with the same four shrinkage methods to get the residuals^1^, a total of 24 methods. The results from `BGLR` are poor and the results from the machine learning methods are relatively independent of the shrinkage method. So, we only show results from our proposed logistic classifier and from the four machine learning methods using the unpre-processed data, a total of eight methods. For each of these, we have five proportions and we only show results from .65 because that was the only case where AEN performed slightly better than the other choices – in accordance with our findings for the real data. For other proportions (except .95) ALASSO was slightly better in simulations.

Suppose we define the two classes by using the 65th percentile and compare the mis-classification rates of the eight methods. These are shown in Fig. 1. Clearly, the best classification method in terms of misclassification error is RFs, a non-sparse method that, by design, can handle nonsparse data such as we have tried to generate. The surprising point is that other non-sparse methods perform considerably worse. The single sparsity method SVMs (with radial basis function kernel) does a bit worse than RFs but better than the other non-sparse methods. In the lower panel, among the doubly sparse methods, SCAD does poorly while the other three perform similarly with AEN doing slightly better. The RAW version with all the variables entering the OPM, gives no sparsity, and does slightly worse. For the 65th percentile, RFs retained around 3,500 of the possible explanatory variables; ALASSO retained low 50s; and AEN retained mid-40s, depending on the run. That is, in terms of sparsity, RFs have none while AEN does noticeably better than ALASSO. Roughly, the best (non-sparse) performance is achieved by RFs, the singly sparse method performs a little worse than RFs, and the doubly sparse methods perform worst, but the best of these, AEN, is nearly as good as RFs. Indeed, if the data is not perfectly sparse, as would be expected here, achieving prediction with a sparse method that is nearly as good as a non-sparse method is generally going to be more useful because it allows interpretability.

**Figure 1:**
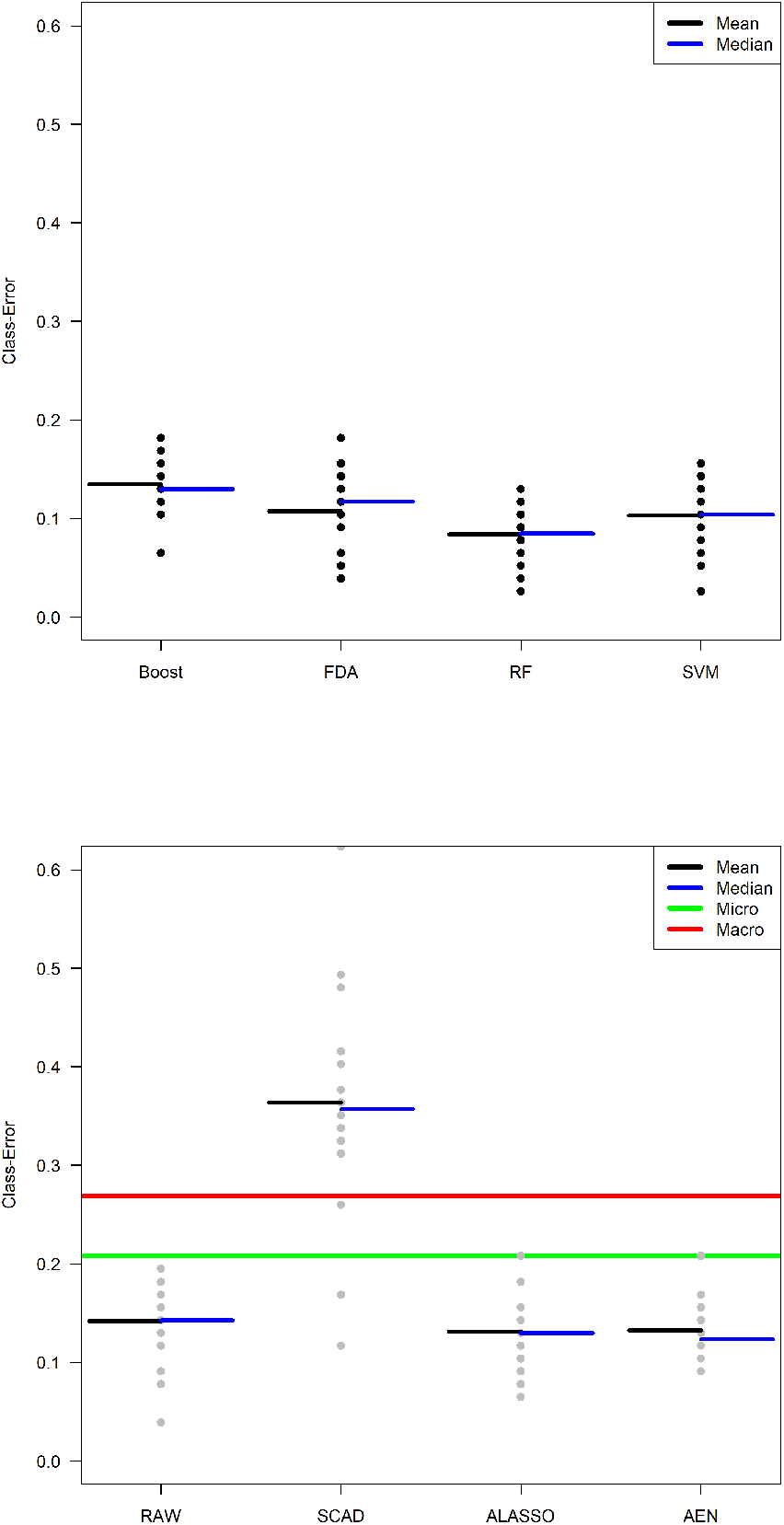
Performance of eight classification methods on the simulated data from Subsec. 4.2. In terms of the misclassification rate, RF is the best, SVM is second, and ALASSO, and AEN are tied for third.

For higher percentiles, the OPM methods tend to perform better relative to RFs, and AEN tends to perform better relative to ALASSO in misclassification but all three methods become more similar in performance and improve slightly. The sparsity level of RFs continues very low while AEN and ALASSO become more similar in terms of sparsity (around 60 variables chosen). Overall, RFs seem to be better for prediction but if the goal includes model identification then AEN-based OPM is the best.

The bottom panel in Fig. 1 shows lines corresponding to misclassification errors using only one type of variables. It is seen that the micro/genomic variables give a lower misclassification error than the macro/phenotypic variables but that both perform worse than using a single data type. In this panel, we used the (standard) BGLR package with a LASSO penalty for the genomic variables and ridge regression for the macro variables. Technically, no penalty is needed for the macro variables, but using ridge regression in this context can only improve the results.

### 4.4 Run-time

To conclude these examples, it’s important to give the running times for computing. All the computations for the OPM were done at The Holland Computing Center (HCC) at the University of Nebraska-Lincoln. The Crane cluster has 548 nodes with 2 Intel Xeon CPU/16 cores per node. The initial data for our simulations were generated from *Synbreed* and took 5-10 minutes.

We present the running times for our real data in two parts. First, we ‘residualize’ the data by preprocessing. That is, we use one of the three shrinkage methods to replace the macro variables by their residuals with respect to the micro-variables as in see (2). Second, we use the pre-processed data in one of five methods – four machine learning methods and three versions of our method (depending on the penalty in (2)). This is reasonable because residualizing the data is a logically separate process from constructing the classifer.

The first four methods could be applied to the unpre-processed data or to the residualized data; the running times for these two cases were essentially the same. Moreover, Table 1 shows these four procedures ran in basically half a minute or less. The last column shows that the running time for OPM ranged from 1.66 to 35 minutes and assumes that the macro variables were already residualized. The time required to residualize the data varied from 10 seconds to 2 minutes. Thus, adding 10 seconds to 2 minutes to each entry in the Table 1 gives the total running time to pre-process the data and then construct the classifier.

**Table 1:**
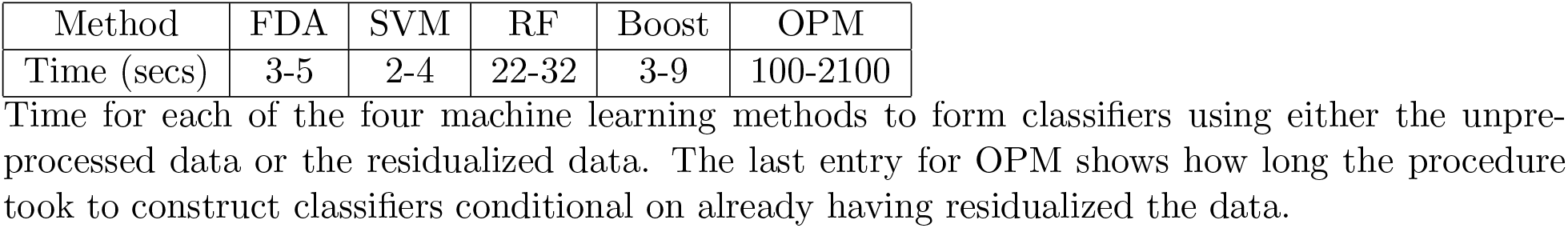
Running times of methods

| Method | FDA | SVM | RF | Boost | OPM |
| --- | --- | --- | --- | --- | --- |
| Time (secs) | 3-5 | 2-4 | 22-32 | 3-9 | 100-2100 |

Time for each of the four machine learning methods to form classifiers using either the unprocessed data or the residualized data. The last entry for OPM shows how long the procedure took to construct classifiers conditional on already having residualized the data.

The wide range for OPM is the result of testing 34 different *λ* values (0.2, 0.4, …, 20) with different individual running times; the smallest values of *λ* took up to 35 minutes. For each one of the 4 methods for including the macro variables (RAW, SCAD, AEN and ALASSO) for each of the 5 quantiles (0.5, 0.65, 0.75, 0.85 and 0.95) 680 (34 * 4 * 5) combinations were considered. Each of these combinations was repeated 20 times, one for each random partition for forming training and testing sets. Hence, the total number of unique combinations using the OPM was 13,600 (680 * 20). For each job, a total of 16 GB ram was allocated for 2 hours.

It is seen that our methods require longer running time than other methods. In general, however, our methods give as good or better results so the improvement in performance may be worth the computing time. Moreover, if our code were optimized, like the packages used for the machine learning methods, likely its running time could be reduced.

## 5. REAL DATA – MAIZE

To see how our methods perform on real data, we consider the maize data recently presented and analyzed (for a different purpose) in Liang et al. (2020). The data we use are available at https://www.sciencedirect.com/science/article/pii/S1674205220300654#mmc2. The original study imputed missing phenotype data using the `Phenix` method described in Dahl et al. (2016) and took other measures to make the data analysis-ready. In particular, even though imputation was used – and could under-represent the complexity of the data – it should affect our methods equally so the comparison will still be fair. For our purposes, this is very high-quality data. There were *n* = 277 data points and we use the same size of training and test sets as given in Subsec. 4.3. From the microvariables in the data set, we select 5,000 at random for 20 different training-testing partitions. We choose five different macroscopic variables (tillering numbers in various environments) to serve as main traits for five analyses. These five are already discrete, assuming values 0, 1, 2, We dichotomized this into five 0-1 main traits by defining *Y* = 0 for zero tillers and *Y* = 1 for one or more. Then from the remaining macrovariables that are continuous, we choose the 4 that are most highly correlated with each of the candidate main traits we dichotomized. That is, a total of 20 highly correlated variables are used in the overall analysis. This gives us five individual analyses we can perform so that the different choices of main trait loosely corresponds to the five percentiles we use in Sec. 4.3. There are other analyses we could have performed on this data using continuous data such as cob length or plant height as the main trait.

Our real data results are broadly similar to the simulation results, at least for proportions around .65, see Fig. 2. RFs remain among the better classification methods but not as clearly so. In addition, the real data analysis indicated more clearly that the doubly sparse method based on AEN is the best choice for model identification and classification outperforming the ML methods – even ‘RAW’ often outperformed the ML methods. Indeed, the top panel of Fig. 2, SVM, FDA, and RFs are nearly the same. with boosting a little worse. In the second panel, AEN is clearly the best of the OPM (and other) methods. In terms of sparsity, SVMs being singly sparse, depends on all the variables as does FDA. RFs used over 4000 variables. Boosting actually gave good sparsity but not as good as AEN. By contrast, for very slight loss (if at all – sometimes a gain) in classification performance, AEN requires only around 45 variables. This sort of behavior is very similar to the behavior of the other four analyses for the other four tillering variables taken as main traits. Overall, the outperformance of AEN compared with the other methods was uniformly higher on real data was than in simulations.

**Figure 2:**
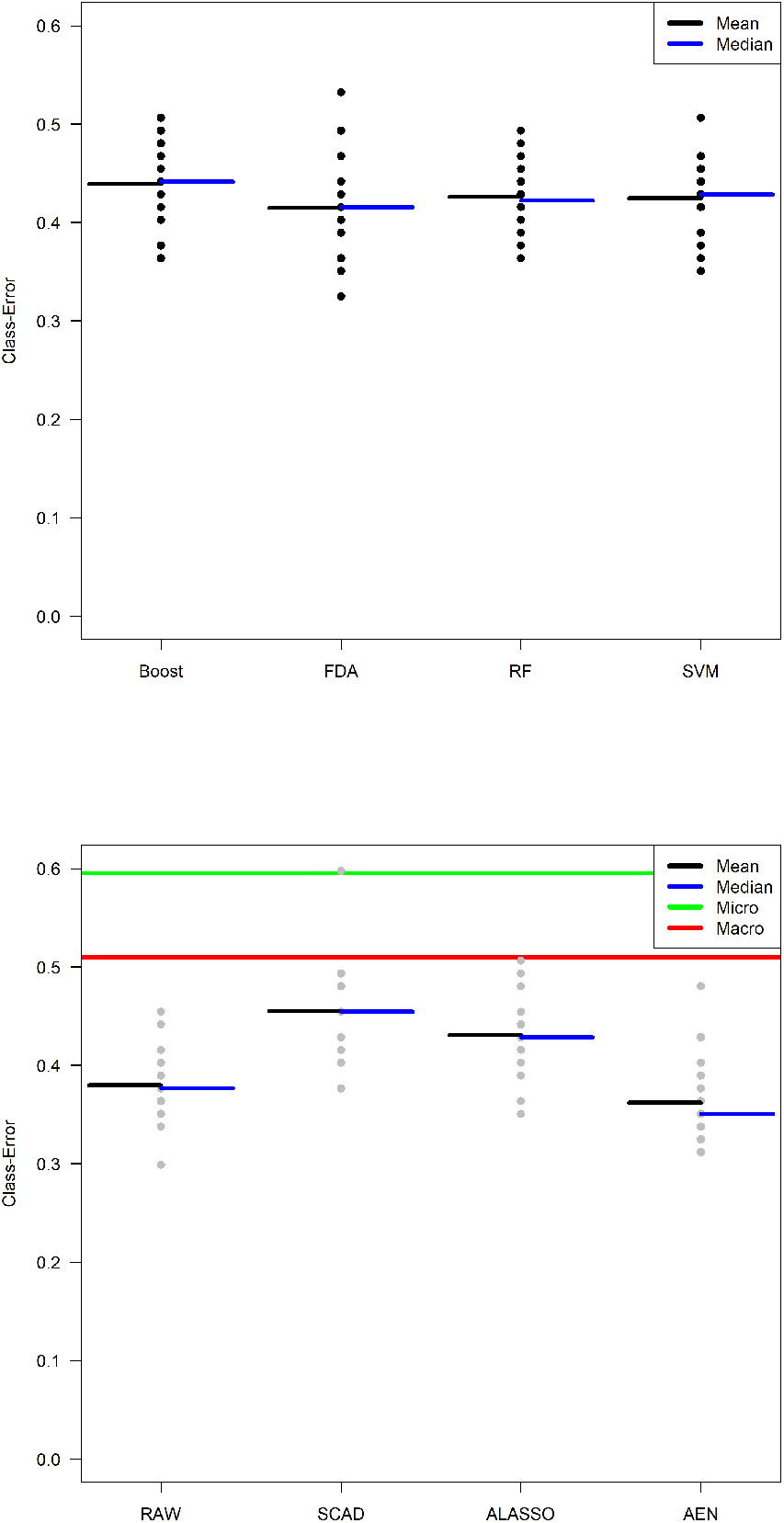
Performance of eight classification methods on Liang et al. (2020)’s data. AEN has the best performance followed by RAW, and the remaining methods are roughly tied. In some cases, the black and blue lines coincided and so cannot be seen separately.

As can be seen in Fig. 2 (and the other real data examples not shown here) the performances of all methods on one of the real data main traits are noticeably worse than on the simulated data.

This is no surprise. Even though the simulations generate relatively complex data, the real data is even more complex. We interpret this to mean that the sparsity built into the simulations, albeit reduced by the complementary QTLs, is still enabling techniques that seek sparsity to give better results. By contrast, in the real data, where sparsity is desired but cannot be safely assumed to the same extent, the techniques continue to try to find what sparsity there is but are necessarily less successful. In this sense, the simulations do not match the generation of real data as closely as we would like despite substantial effort at modeling. On the other hand, the qualitative similarities are reasonably strong and that is enough for present purposes.

The bottom panel in Fig. 2 shows lines corresponding to misclassification errors using one type of variables. It is seen that the micro/genomic variables give a lower misclassification error than the macro/phenotypic variables but that both perform worse than using a single data type. In this panel, we used the (standard) BGLR package with a LASSO penalty for the genomic variables and ridge regression for the macro variables. Technically, no penalty is needed for the macro variables variables, but using ridge regression in this context can only improve the results.

We find that the variables forming the best classifiers for the five main traits we examined were always phenotypic variables representing some aspect of tasseling or ‘number of tillering plants’ meaning a direct count of the number of plants in a row of a particular genotype that are producing tillers. So, we find an association between tillering of individual plants with how prevalent tillering is and the growth of the male part of the plant (the tassel). It is tempting to conjecture that some genes or pathways may be conserved between regulating the development of vegetative branches (tillers) and inflorescence branches (the branches on tassels) so that a line with more tillers might also have more tassel branches and vice versa.

## 6. REAL DATA – SOYBEAN

Here we demonstrate the performance our technique on another data set called `SOYNAM`. This data set was presented and analyzed using conventional methods in Diers, Specht, Rainey, Cregan, Song, Ramasubramanian, Graef, Nelson, Schapaugh, Wang, Shannon, McHale, Kantartzi, Xavier, Mian, Stupar, Michno, Charles, Goettel, Ward, Fox, Lipka, Hyten, Cary, and Beavis (Diers et al.) and Xavier, Jarquin, Howard, Ramasubramanian, Specht, Graef, Beavis, Diers, Song, Cregan, Nelson, Mian, Shannon, McHale, Wang, Schapaugh, Lorenz, Xu, Muir, and Rainey (Xavier et al.) who also provide a detailed description of it. Briefly, the SoyNAM project is comprised of 40 biparental populations (140 individuals per population) sharing a common hub parent (IA3023). The hub parent was crossed with elite parents (17), plant introductions (8), and parents with exotic ancestry (15). The resulting 5,600 (40×140) accessions were observed in 18 location×year combinations (environments). Also, genotypes were scored for 9 traits: lodging ([1-5], 1 with almost all plants erect, 5 all plants down), plant height (cms), days to maturity (R8), grain yield (y), moisture, (m) protein content (%, pro), oil content (%), fiber (fi), and seed-size.

The main trait of interest here is lodging. It is important because it can reduce photosynthesis and hence yield by as much as 10% with consequent effects on profitability. In this study, values greater than 2.5 (3, 3.5, 4, 4.5, and 5) were represented as 1 and values below them (1, 1.5, 2, and 2.5) were represented as zero. In effect, we dichotomized a discrete variable resulting in 2,402 genotypes in group 0 and 2,626 in group 1.

The remaining eight traits were treated as macroscopic variables. For consistency with our earlier work, we completed the eight traits to a set of 20 macroscopic variables by adding 12 interaction terms; namely, height× R8, height×yield, R8×yield, R8×moisture, yield×moisture, yield×protein, moisture×protein, moisture×oil, protein×oil, protein×fiber, oil×fiber, and oil×seed-size. Adding these interaction terms makes the set of macroscopic variables more informative.

One of the interesting features of this data is that unlike many data sets with a large number of genomic/micro variables, the macro variables seem to have a substantial majority of the information about the main trait in them. That is, if we take the main trait as lodging then most of the variability in lodging is explained by yield. We comment that it is more typical to choose yield and as the main trait and use lodging as a secondary trait to predict it. Here, we reverse this because lodging is a categorical variable that is worth studying in its own right. The high degree of correlation between Lodging and the other macro variables can be seen in Table 2.

**Table 2:**
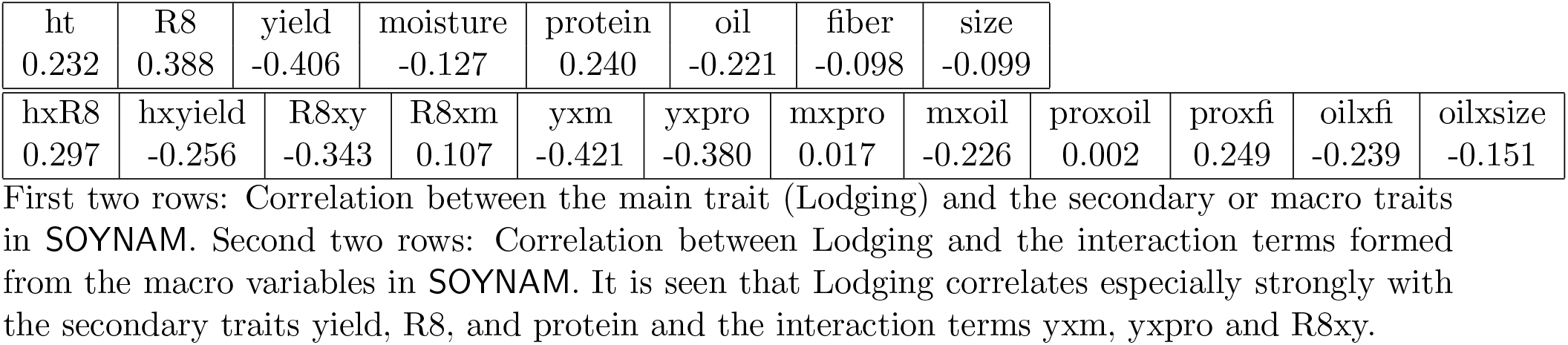
Correlations of Lodging with the macro traits

|  |  |  |  |  |  |  |  |
| --- | --- | --- | --- | --- | --- | --- | --- |
| ht | R8 | yield | moisture | protein | oil | fiber | size |
| 0.232 | 0.388 | -0.406 | -0.127 | 0.240 | -0.221 | -0.098 | -0.099 |

|  |  |  |  |  |  |  |  |  |  |  |  |
| --- | --- | --- | --- | --- | --- | --- | --- | --- | --- | --- | --- |
| hxR8 | hxyield | R8xy | R8xm | yxm | yxpro | mxpro | mxoil | proxoil | proxfi | oilxfi | oilxsize |
| 0.297 | -0.256 | -0.343 | 0.107 | -0.421 | -0.380 | 0.017 | -0.226 | 0.002 | 0.249 | -0.239 | -0.151 |

First two rows: Correlation between the main trait (Lodging) and the secondary or macro traits in SOYNAM. Second two rows: Correlation between Lodging and the interaction terms formed from the macro variables in SOYNAM. It is seen that Lodging correlates especially strongly with the secondary traits yield, R8, and protein and the interaction terms yxm, yxpro and R8xy.

The RIL genotypes were sequenced with a 6k array delivering a total of 5,300 maker SNPs. After applying conventional quality control, e.g., discarding those SNP’s with more than 20% missing values, a minor allele frequency smaller than 0.03, etc., we had 4,325 microscopic genomic variables for analysis. Again, for consistency with our earlier work we produced a set of 5,000 markers by choosing a random sample of 675 artificial and independent SNP’s to add and to the actual set of SNP’s.

Here, we only analyzed data from the NE 2012 location. After alignment (phenotypic and marker data) and keeping only genotypes with phenotypic information for all 9 traits, 5,028 genotypes remained for analyses. We then selected 6 random samples (without replacement) of size 277 from the 5,028 genotypes. For each sample we used 20 repetitions of our method essentially bootstrapping the data. This mimics the case that we have one real data set that is too small to be divided.

The misclassification errors for the four ML methods, three doubly penalized methods, and the OPM method (with only one penalization to obtain the residuals) are given in Fig. 3 following the same conventions as Fig. 2. It is obvious that the two best methods are RAW and AEN and for all practical purposes they tie predictively. To look into what these two methods are doing ‘behind the scenes’ we counted how many of the macro- and micro-variables each included. Curiously, both led to about the same sparsity, choosing, on average, around 10-11 of the macroscopic variables, see Table 3. Indeed, RAW chose none of the micro-variables and AEN only chose one or two in a few runs. Both methods successfully eliminated the 675 decoy variables. This illustrates that even though the ‘typical’ case is that high dimensional data crowds out low dimensional data the reverse can happen too: In this case, the macroscopic variables RAW in and the adjusted macroscopic variables in AEN crowd out the microscopic variables, perhaps because we included interaction terms.

**Figure 3:**
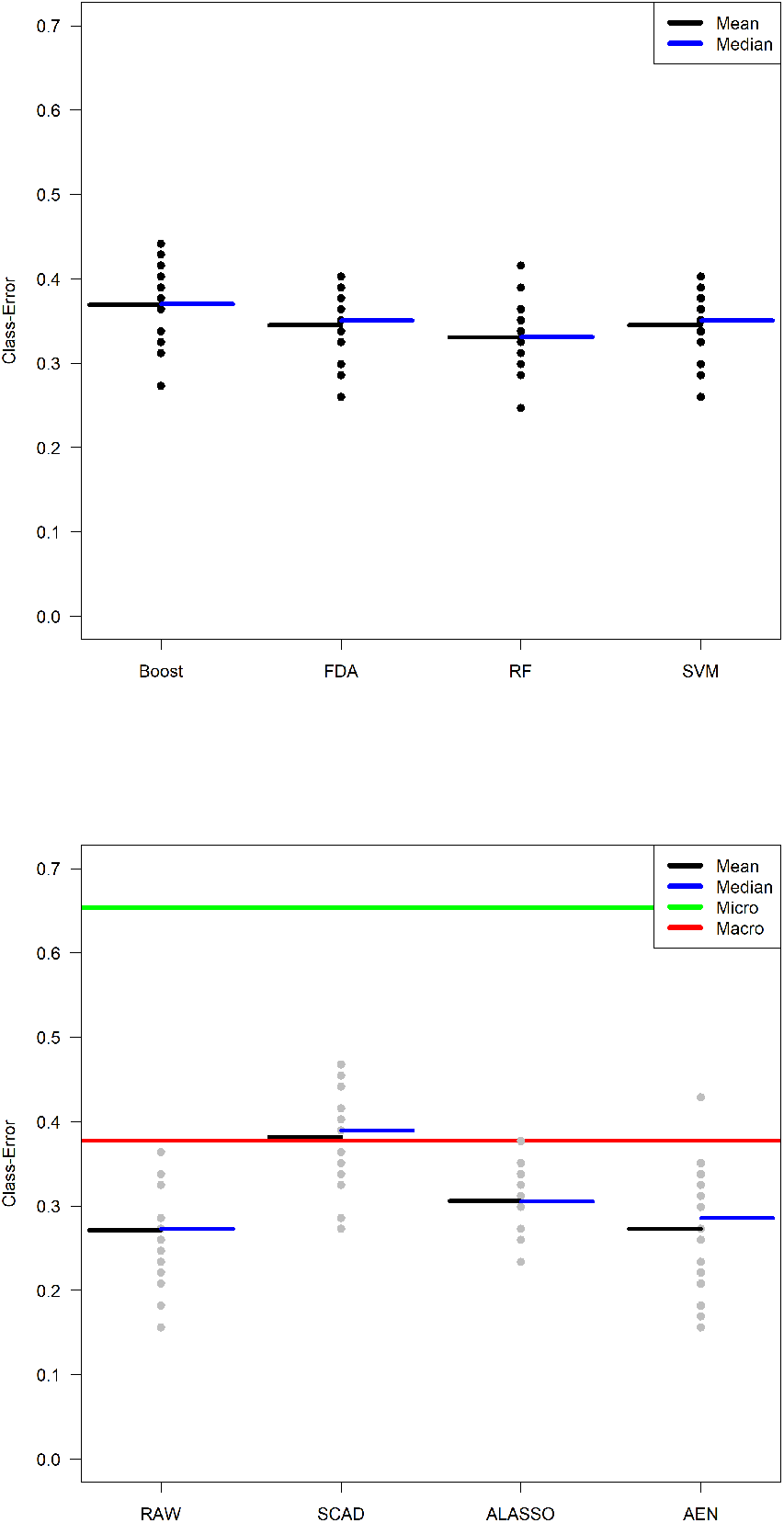
Performance of eight classification methods on the modified soybean data. AEN and RAW tie for the best performance. Note that BGLR, used for only one class of variables give worse performances than the sparsity based methods – even if one reverses the classification for the micro variables.

**Table 3:** No. of Macro Traits Selected

| Proportion | .5 | .65 | .75 | .85 | .95 |
| --- | --- | --- | --- | --- | --- |
| RAW | 11.75 | 12.1 | 11.75 | 10.6 | 9.75 |
| AEN | 12.4 | 11.7 | 11.9 | 10.95 | 10.5 |

Number of secondary traits (macroscopic variables) chosen by RAW (OPM using LASS) and AEN (AEN followed by OPM using AEN) for five different proportions.

As a further comparison of the two methods, RAW and OPM+AEN, we found the SE’s of their misclassifiaciton rates, see Table 4. It is seen that the differences are small, only in the second decimal place; the biggest difference is .011 (for .65) a little over 1% or around *±*.02 in the sense of a 95% confidence interval for the mean misclassifiaction error. We interpret this to mean that RAW and AEN are tied, but the tie is slightly in favor of RAW for the purposes of classification.

**Table 4:**
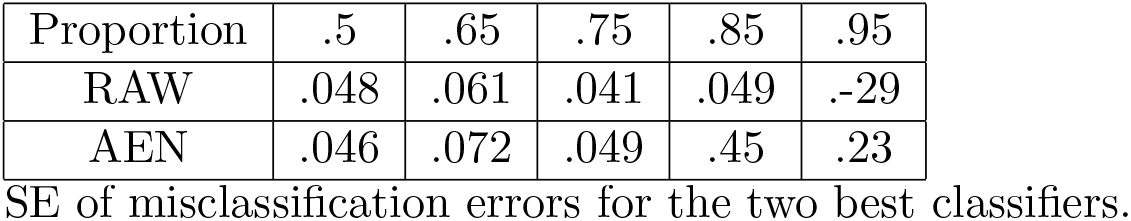
SE’s of OPM + shrinkage

**Table 5:** Abbreviations

SE of misclassification errors for the two best classifiers.
| Table 5: Abbreviations |  |
| --- | --- |
| Abbreviation | Meaning |
| AEN | adaptive elastic net |
| ALASSO | adaptive LASSO |
| Boost | boosting |
| CV | cross-validation |
| FDA | Fisher's discriminant analysis |
| HCC | Holland Computing Center |
| LASSO | least absolute selection and shrinkage operator |
| LD | linkage disequilibrium |
| ML | machine learning |
| NR | Newton-Raphson |
| OPM | one pass method |
| PLL | penalized log-likelihood |
| QTL | quantitative trait locus |
| QTLc | complement of a set of QTL's |
| RBF | radial basis function |
| RF | random forest |
| RIL | recombinant inbred lines |
| RR | ridge regression |
| SCAD | smoothly clipped absolute deviation |
| SNP | single nucleotide polymorphism |
| SVM | support vector machine |
| Var | variance |

The net result of this analysis is that the RAW and the AEN methods are predictively equivalent once all sources of error are taken into account.

However, recall that when we use the AEN penalty, the macro variables are first adjusted by sparsely regressing them on the micro variables with LASSO. Then, the residuals are put into the OPM with the AEN penalty. Thus, when the AEN method selects variables only from the adjusted macro variables some micro variables are included because of the adjustment. By contrast, with RAW, only the non-adjusted macro variables are included. Thus, if it is sincerely believed than none of the micro variables have any relation to the main trait, RAW gives ever-so-slightly better results. On the other hand, if it is believed that at least some of the micro variables are related to the main trait then even the small loss of classification accuracy is probably worth identifying them via the adjustment to the macro variables. In this sense, the modeling provided by OPM with AEN is better than by RAW. In this case, our interpretation is that RAW and AEN are tied, but the tie is actually in favor of AEN since the RAW model is implausible.

This situation is really only possible when the correlation between the main trait and secondary traits is sufficiently high but shows that macro variables (adjusted or not) can crowd out the microvariables. Thus, even in this case, for which our method was not explicitly designed, the cost in terms of misclassification is very small indeed and still provides a selection from both sets of variables (macro and micro) that can in principle be interpreted.

We comment that an intermediate case – using no penalty in place of AEN for instance – will not in general provide viable solutions. The OPM, although strong as a computational method, does not by itself ensure consistency of variable selection or parameter estimation.

## 7. DISCUSSION

The classification methods we consider in our study are not exhaustive, as there are numerous such methods. As a related point, it is possible to get double sparsity from singly sparse methods. The basic technique is similar to out-of-bag error estimates for RFs: permute (or just leave out) values of variables and see whether the error increases much. The problem with this approach is its ability to eliminate excess variables reliably is not very well explored.

We believe that the present analysis, although well-motivated, will likely be limited to specific data problems for instance those that have some form of sparsity. Another way to say this is that the data probably cannot deviate too much from the examples computed here.

One reason the OPM may work well is the obvious one: Even in high dimensions, the actual function may be locally well-approximated by a linear function, and, on planes, it is reasonable to optimize locally one variable at a time.

We have argued that starting with residuals from macro variables gives better results than the other approaches we tested. We suggest the reason for this is the obvious one, namely, that if the residuals from the macroscopic variables are given ‘first crack’ at being in the classifier then some of them will be retained. Thus, the later (microscopic) variables are disfavored unless they are really useful.

For the mid-range of proportions (that were of most concern to us) AEN emerged as the recommended shrinkage method amongst those we considered. A heuristic explanation is that ALASSO does shrinkage and variable selection in one step. As such it is designed to give model sparsity and this may explain its marginally better performance in simulations in terms of misclassification for proportions between .75 and .95. In results not shown here, ALASSO often gave fewer variables if not by much. On the other hand, ridge regression (RR) does not in general give sparsity. The AEN penalty is a weighted combination of the RR penalty and the LASSO penalty. Both of our real data sets and our simulated data had sparsity, sometimes moderate and sometimes high. Accordingly a method like AEN that pushes to sparsity but not excessively turns out to do best. Otherwise put, the adaptivity of AEN to the sparsity level in the data may be what allows the AEN to give generally competitive if not superior results.

In the bottom panels of Fig. 1, 2, 3 we did not simply modify our software for the single variable type lines. If we had, Fig. 1 and 3 would have been qualitatively unchanged. That is, in simulations or ith `SYNAM`, with the exception of the SCAD penalty, our method always had a comparable or lower classification error using the two data types than using either of the single data types. However, with maize data the comparison between using both data types and single data types was more nuanced. It depended on extra features of the data that were not considered in our generic method, such as collinearity among explanatory variables and similarity of the explanatory variables *physically* to the response *Y*. Accordingly, in both figures we compared our method against the standard methods used for single data types as indicated. Moreover, in our exploratory work (not shown here) we have been led to conjecture that the performance of our method also depends on the complexity of the trait *Y*. Echoing the work of Jeong et al. (2020), this is difficult to quantify, but intuitively, the more complex *Y* is, the more our method using both data types should outperform our method restricted to a single data type. This is reasonable because more complex traits require more data to explain them.

Four more technical points to note. First, our method can be generalized to three or more data types. Sparsely regress one data type on another, repeat with the residuals to get a second set of residuals. These represent the data only in the first data type. Then use our method here on the remaining two data types and apply the OPM letting the smallest dimensional data type enter first. Second, we have only used binary classification here, but the method as written extends to multiclass classifiers in the obvious way. Third, our analysis and conclusions have included sparsity as well as prediction because we have taken the research goal as including model building so that sparsity is essential for determining which variables are important. If the goal were simply finding a good class predictor then sparsity would be less relevant and we would have been led, likely, to RFs as the overall best choice even though in some instances other methods, including ours, performed better. This is an active area of research and we would hesitate to make any definitive statement. Four, we have used tillering numbers or lodging as the main traits in our real data examples. The consensus seems to be that too many tillers is bad for yield but the question of their effect and genetic control remains of interest; see Zhang et al. (2019) for a recent overview. Likewise, lodging can also be costly. We do not address these details because our goal has been to produce a good classifier for complex main traits. We can, however, conjecture that tillering responds primarily to the presence of tassels because the prevalence of nearby tillers (in the same genotype) may simply also be the result of tasseling. At this point, however, we are not able to offer a convincing agronomic interpretation from our analysis of the `SOYNAM` data apart from suggesting that some of the micro variables must be related to lodging, possibly mediated by one or more of the macro variables.

## 8. CONCLUSION

Our goal here was to give a principled method that could be used to generate classifiers when two types of explanatory variables are available for each subject. The two data types were regarded as differing in dimension, one being high-dimensional (microscopic) and the other being low-dimensional (macroscopic). It was also assumed that some of the variables in the low-dimensional data were essential to forming a good classifier so our method would not let the high-dimensional data (that perhaps did not contribute as much information, at least per variable) swamp the entirety of the low dimensional data. Thus, our method – a sparse logistic classifier – is intended for complex responses (main traits), where using more than one data-type is essential to modeling. The technique we have advocated is to construct a sparse logistic classifier by sparsely regressing the macroscopic variables on the microscopic variables, and using the residuals to represent the information in the macroscopic variables not in the microscopic variables. We did this sparse regression in a variety of ways finding that LASSO gave the best results. Then, to find the logistic classifier, we used an AEN penalty and a one-pass-method so that the explanatory variables and their coefficients coefficients could be effectively computed.

We compared our method with variants on it and with several standard machine learning classifiers. Overall, our analyses strongly suggest that for the type of classification problems we have considered, the combination of sparse regression with LASSO, priority of macro residuals, and OPM with AEN to find the coefficients in a penalized logistic regression gives the best results for the joint purposes of prediction and model identification.

## AVAILABILITY OF DATA AND MATERIALS

All code used for the computations presented here is available at https://github.com/royarkaprava/OPM.

## ACKNOWLEDGEMENTS

The authors thank Holland Computing Center for providing computing resources and Zhikai Liang & James Schnable for making their data and insights accessible to us.

## Footnotes

1 Actually, we also tested RVMs with an RBF kernel and SVM with a Laplace kernel. The available code for the former often would not run and the latter gave very poor results. This amounted to testing an extra eight methods.

